# Integrative single-cell analysis by transcriptional and epigenetic states in human adult brain

**DOI:** 10.1101/128520

**Authors:** Blue B. Lake, Song Chen, Brandon C. Sos, Jean Fan, Yun Yung, Gwendolyn E. Kaeser, Thu E. Duong, Derek Gao, Jerold Chun, Peter Kharchenko, Kun Zhang

**Author notes:** Equally contributed authors (Blue B. Lake, Song Chen, Brandon C. Sos, Jean Fan). Corresponding Authors: Kun Zhang; Peter Kharchenko; Jerold Chun.

## Abstract

Detailed characterization of the cell types comprising the highly complex human brain is essential to understanding its function. Such tasks require highly scalable experimental approaches to examine different aspects of the molecular state of individual cells, as well as the computational integration to produce unified cell state annotations. Here we report the development of two highly scalable methods (snDrop-Seq and scTHS-Seq), that we have used to acquire nuclear transcriptome and DNA accessibility maps for thousands of single cells from the human adult visual and frontal cortex. This has led to the best-resolved human neuronal subtypes to date, identification of a majority of the non-neuronal cell types, as well as the cell-type specific nuclear transcriptome and DNA accessibility maps. Integrative analysis allowed us to identify transcription factors and regulatory elements shaping the state of different brain cell types, and to map genetic risk factors of human brain common diseases to specific pathogenic cell types and subtypes.

## Introduction

Our brain is an enormously complex network consisting of 100 billion spatially organized and functionally connected neurons embedded in an even larger population of glia and non-neural cells. Producing a complete cell atlas of the human brain would require highly scalable and unbiased approaches. Recent advances in droplet-based technologies has greatly enhanced the throughput of single-cell RNA-sequencing (RNA-seq)^1-3^, enabling simultaneous transcriptomic profiling on the order of tens of thousands of single cells. While these methodologies limit depth of coverage, they enable extensive cell type and state classification, providing unique expression signatures to resolve functional heterogeneity existing within tissues. However, a reliance on live intact cells impedes the goal of generating comprehensive maps of transcriptionally defined cell states for all tissues and organs that would be needed to build a complete cell atlas of the human body. Many complex tissues are not amenable to cellular dissociation nor are they readily available from fresh biopsies. Nuclear isolates are readily accessible from fresh or achieved tissues and provide sufficient RNA for accurate prediction of cellular expression levels^4-7^ without artefactual signatures associated with tissue dissociation^8^, and are highly amenable to single-cell genomic studies ^9, 10^. We have recently demonstrated that Single-nucleus transcriptome Sequencing (SNS) can resolve neuronal subtype diversity across multiple human cortical brain regions ^7^, at a relative high sequencing depth (∼8 million reads per cell). However, scaling these studies was limited by the throughput (up to 96 cells per microfluidic chip) and high cost. In addition, non-neuronal nuclei are smaller and difficult to capture with Fluidigm C1 microfluidic chips. Therefore, there is not only a growing need for more efficient methods for nuclear RNA sequencing of various cell types in human archived tissues, but also for co-profiling genomic attributes, such as epigenetic state, in order to build a more comprehensive picture of a cell’s overall phenotypic potential. While methods for characterizing chromatin accessibility have been reported^11-13^, they remain to be demonstrated on highly heterogeneous archived human tissues, like the brain, at scale. Here we report two highly scalable methods for quantifying nuclear transcripts and measuring DNA accessibility at the single-cell level on human archived tissues, and have applied both methods to the first integrative analysis of single-nucleus transcriptomes and chromatin maps of human tissues.

## Results

To overcome the limitations associated with single-nucleus RNA sequencing, we have adapted a droplet-based methodology^2^ to analyze single nuclei, termed snDrop-seq, that we find can provide a higher-scale assessment of neuronal and non-neuronal diversity in human post-mortem brain samples. The application of this methodology to isolated nuclei necessitated certain modifications (see **Methods**) that included coating all inner surfaces with bovine serum albumen (BSA) to prevent non-specific binding by nuclei, and heat treatment of droplet-encapsulated nuclei to ensure complete lysis without introducing excessive RNA degradation (**Fig. S1a**). Subsequently, we applied the snDrop-seq pipeline (**Fig. 1A**) to both the human adult visual cortex (Brodmann Area 17 (BA17) or V1) and frontal cortex samples (BA10 and BA6) from four different individuals (**Table S1**). We generated 16,262 visual cortex singlenucleus data sets and 9,794 frontal cortex data sets, of which 15,819 and 8,755, respectively, were resolved into neuronal and non-neuronal cell types (**Fig. 1B-E**, **Table S2**). These libraries were sequenced to a median of 5,247 usable reads (2,483 for frontal cortex), with the majority of mapped reads falling within intronic regions (**Fig. S1D-E**), and predominantly to the 3’ ends of transcripts (**Fig. S1F**), consistent with poly-A capture and 3’-end counting for not only mRNA transcripts, but also the pre-mRNA abundant in nuclei^14^. In comparison with other RNA-seq methodologies (**Fig. S2**), both snDrop-seq and scDrop-seq^2^ methods showed highly comparable, albeit lower, median UMI counts and gene detection rates. However, nuclear data was slightly biased for longer genes (**Fig**. **S2Q-R**), likely reflecting differential transcript processing and export rates associated with genic length and intron fraction^14^. Overall, we detected a median of 1,072 unique transcripts and 826 genes per visual cortex nucleus (**Fig. S2)** and 7,422 total genes detected on average per cell type **(Table S2**). Analysis of transcriptional heterogeneity (see **Methods**) resolved 30 distinct cellular clusters representing 10 excitatory (**Ex**) and 14 inhibitory (**In**) neuronal subtypes, and six non-neuronal cells, including: endothelial cells (**End**), smooth muscle cells or pericytes (**Per**), astrocytes (**Ast**), oligodendrocytes (**Oli**) and their precursor cells (**OPCs**) and microglia (**Mic**) (**Fig. 1B, Fig. S3**). These subpopulations showed cell-type or subtype specific expression profiles (**Fig. 1C, Table S3**), expected marker genes (**Fig. 1D**), and were highly consistent with pooled cell populations from the mouse^15^ and human (temporal lobe)^16^ cerebral cortex (**Fig. 1F**). Comparison with single-cell data generated from the mouse visual cortex^17^ and human temporal lobe^18^ confirmed broad cell type classification and the consistency between nuclear and whole cell data (**Fig. 1G**). Neuronal clusters were annotated based on their correlation values with subtypes previously identified from SNS in six cortical regions^7^ (**Fig. 1H**). In addition to the high correspondence, snDrop-seq permitted finer resolution into sub-populations (e.g. **Ex1** to **Ex1a,b** of the visual cortex) while showing little representation from subtypes not previously observed in these regions (e.g. rostral-specific **Ex2** found only in frontal cortex and caudal-specific **Ex3** found only in visual cortex)^7^. Otherwise, subtypes resolved were found to be highly consistent between these two cortical regions (**Fig. 1I**). This demonstrates the high accuracy of snDrop-seq in resolving neuronal subtype diversity in the cerebral cortex though profiling a larger cell cohort compared with our previous SNS efforts, albeit with ∼1500-fold lower per cell sequencing depth.

**Fig. 1.**
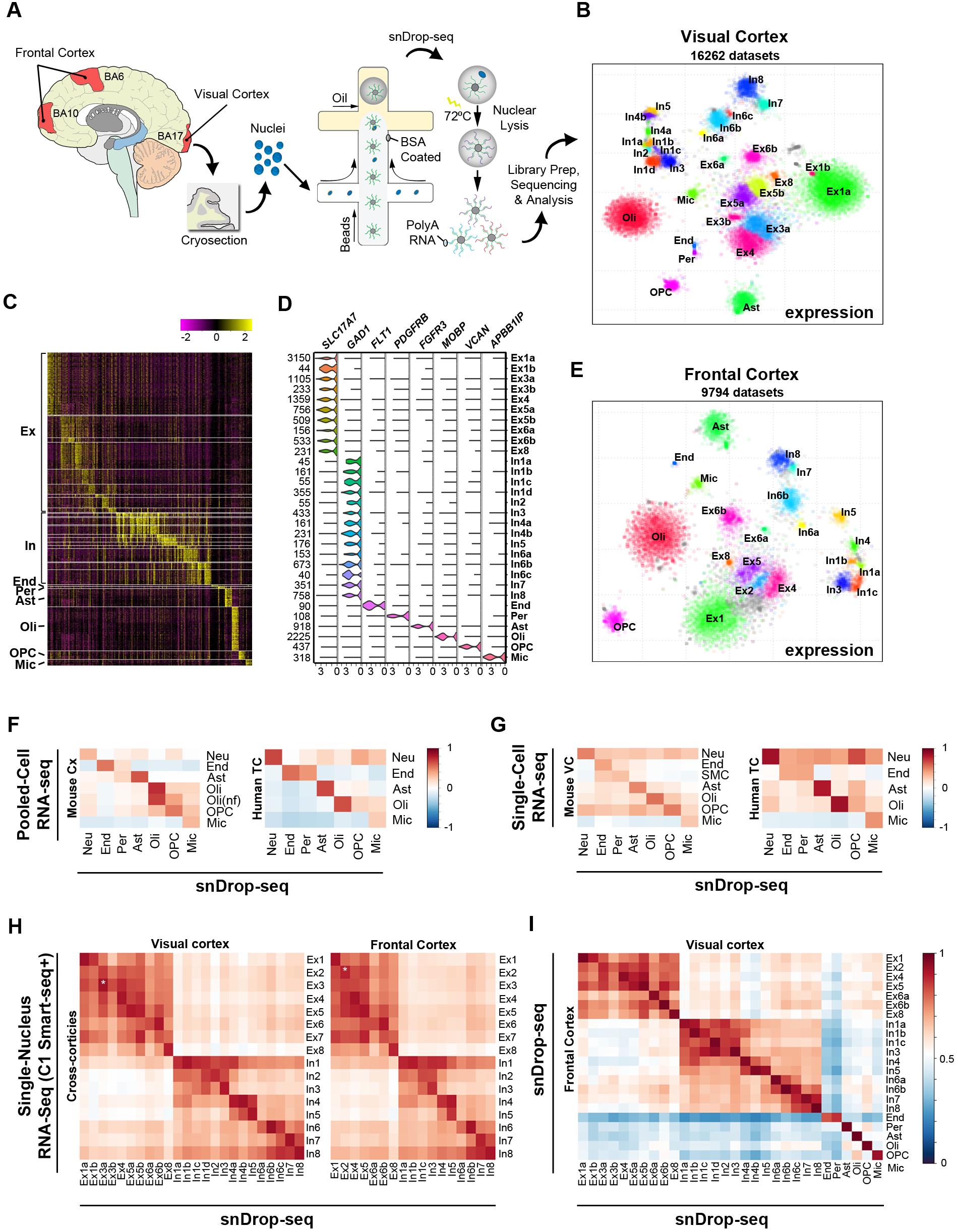
Transcriptional profiling with snDrop-seq resolves major cell types of the visual and frontal cortices. **A.** Overview of single nucleus isolation from the visual cortex (occipital lobe, BA17) and frontal cortex (BA6 and BA10) and subsequent snDrop-seq procedure. **B.** Single nucleus visual cortex data sets (16,262) showed distinct clustering, visualized here using t-distributed Stochastic Neighbor Embedding (t-SNE). Unannotated data sets are shown in gray. **C.** Heatmap of expression of top differentially expressed genes identified across cell types and subtypes of the visual cortex (Table S3). **D.** Violin plots of expression values for type-specific marker genes for the visual cortex. Number of data sets contributing to each cluster is listed. **E.** Distinct clustering of the frontal cortex data sets using t-SNE. Unannotated data sets are shown in gray. **F.** Correlation heatmaps comparing averaged visual cortex snDrop-seq data with average expression values from RNA-seq data from: mouse pooled cortical cell types (*12*) (left, nf = newly formed); human pooled temporal lobe cell types (*13*) (right). **G.** Correlation heatmaps comparing average visual cortex data with average expression values from single-cell RNA-seq data from the mouse visual cortex (*14*) (left) and human temporal lobe (*15*) (right). **H.** Correlation heatmap of snDrop-seq-identified neuronal subtypes (visual cortex or frontal cortex) compared with subtypes previously identified using the C1 single nucleus Smart-seq+ pipeline (SNS, across cortical regions) (*7*). Star indicates distinct region-specific Ex2 and Ex3 sub-populations. **I.** Correlation of average expression values for cell types and subtypes resolved from the visual and frontal cortices.

Excitatory neurons of the visual cortex marked by expression of *SLC17A7* (**Fig. 1D**) were resolved into 10 distinct subtypes, showing enriched marker gene profiles (**Fig. S4A, Table S4**) and that could be distinguished by their spatial orientation within the cortex^19^ (**Fig. 2A, B**). In addition, we identified specific subpopulations located within cortical layers, including the *CARTPT^+^RASGRF2^+^* **Ex1b** of upper layer 2; the distinct *HS3ST5^+^PCP4^-^* (**Ex5b**) and *HTR2C^+^PCP4^+^TLE4^+^* (**Ex6a**) subpopulations in layer 5, the latter bordering on a *HTR2C^-^TLE4^+^* (**Ex6b**) layer 6 population; and the *NR4A2^+^SYNPR^+^* **Ex8** subpopulation in layer 6b (**Fig. 2B**). Inhibitory neurons of the VC, marked by shared expression of *GAD1* (**Fig. 1D**), were resolved into 14 subtypes showing enriched marker gene expression (**Fig. S4B, Table S5**), distinct profiles of canonical interneuron markers (e.g. *VIP, RELN, PVALB, SST*) as well as sub-type restricted expression (e.g. *SHISA8, CA3, CA8*) (**Fig. 2C**). We were further able to resolve spatially distinct inhibitory neuron subpopulations, including: layer 1 *RELN^+^CCK^+^CNR1^+^* **In1a**; upper layer *VIP^+^CALB2^+^TAC3^+^* **In1d**; *PVALB^+^CA8^+^* **In6a** concentrated around layer 4, as well as the more peripheral *PVALB^+^TAC1^+^* **In6b**; and two distinct SST positive subtypes, including the upper layer *SST^+^CALB1^+^* (**In7**) and lower layer *SST^+^CALB1^-^* (**In8**) subpopulations (**Fig. 2D**). Furthermore, given the strand-specificity of snDrop-Seq, we characterized the expression patterns of natural antisense transcripts (NATs), which was not possible with SNS^7^. We identified 26 NATs to be differentially expressed across cell clusters (**Table S7, Fig. S5**), including inhibitory neuron-specific expression of DLX6-AS1 potentially involved in interneuron specification during development^20, 21^ (**Fig. S5B**). Furthermore, we detect a higher proportion of neuronal data sets in the visual over frontal cortex consistent with a higher neuronal density in this region (**Fig. 2E**)^22^. Cell type proportions detected by snDrop-seq were somewhat consistent with those from single-cell analyses on the temporal cortex, with the exception of astrocytes and endothelial cells which appear under-represented. This may reflect a bias in sample processing or a potential limited detection based on lower transcript levels for these cell types (**Fig. S3L-O**). Therefore, snDrop-seq provides not only a more comprehensive cellular profile of human postmortem tissues (**Fig. 2E**), but also insights into region-specific cell-type proportion differences and strand-specific transcriptomic dynamics, attributes not achievable using previous methodologies^7^.

**Fig. 2.**
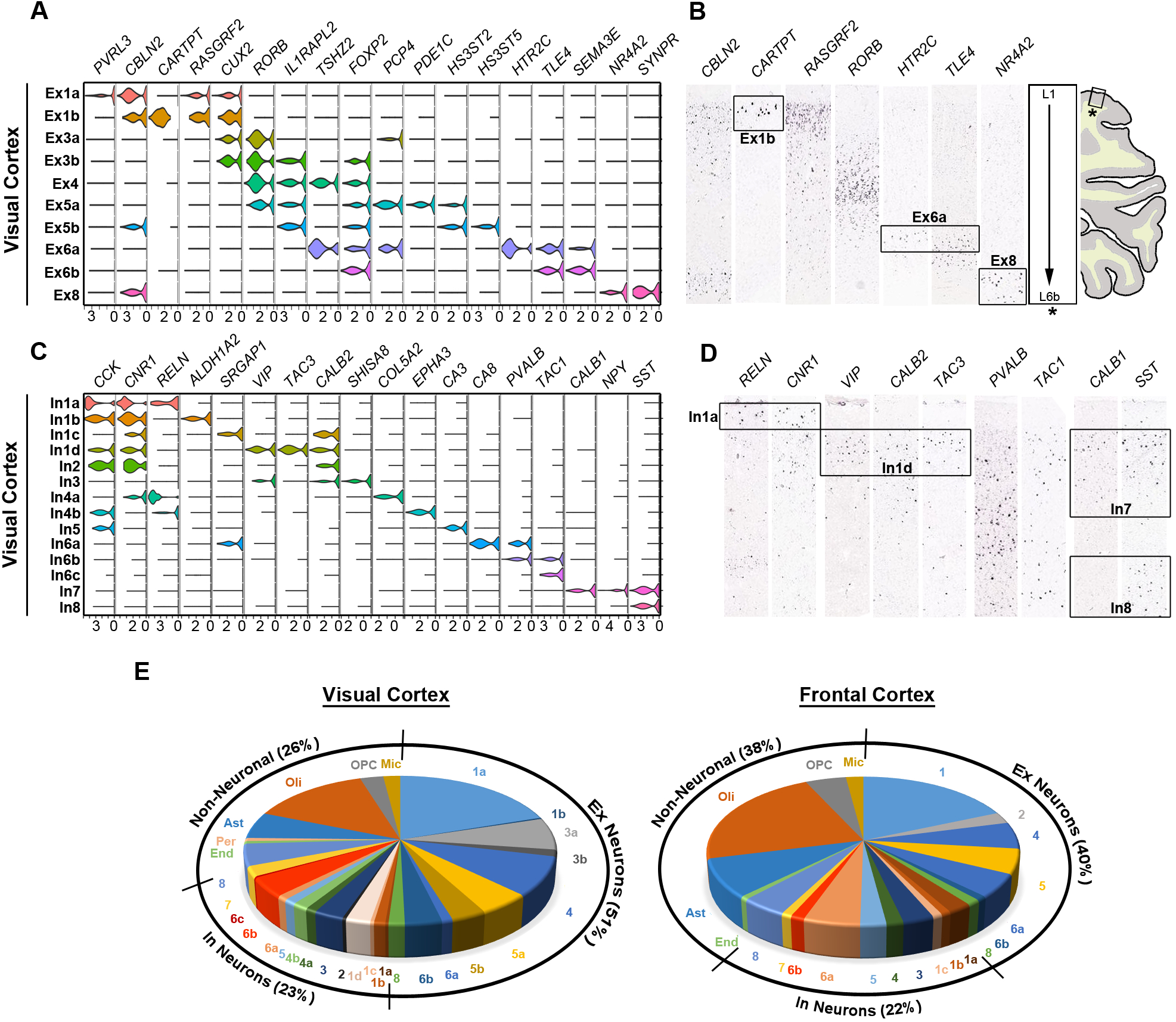
snDrop-seq identifies molecularly and spatially distinct neuronal subtypes. **A.** Violin plots showing gene expression values of layer specific (*7, 16*) and subtype-enriched markers for excitatory neuronal subtypes. **B.** RNA *in situ* hybridization (ISH) stains (Table S6) of the visual cortex for select marker genes shown in (**A**). **C.** Violin plots showing expression values for classical interneuron marker genes (*7*) and subtype-enriched transcripts. **D.** RNA ISH stains (Table S6) of the visual cortex showing select stains for markers in (**C**) and predicted spatial distribution of associated inhibitory neuron subtypes. **E.** Proportion of cell types and subtypes detected from snDrop-seq interrogation of the human visual cortex compared with the frontal cortex.

The epigenetic state of a cell can provide valuable information about its overall phenotypic potential, and reveal gene regulatory processes responsible for establishing and maintaining its function. To investigate the epigenetic configuration of different cells in the human adult cortex at scale, we developed a single-cell DNA accessibility assay by integrating THS-Seq^23^, which uses *in vitro* transcription and an engineered super-mutant of Tn5 transposase^24^ to achieve higher sensitivity and better coverage of distal enhancers than ATAC-seq^25^, with combinatorial cellular indexing^11^ based on a set of 384 customized barcoded transposomes (**Fig. 3A, Fig. S6**). scTHS-seq, confirmed to have a low doublet rate and generating high quality data (**Fig. S7-8, Table S8**), identified 287,381 peaks associated with DNA accessibility regions in combined data sets, covering 144 million base pairs with unique genomic alignments, at the clonal rate of ∼60% and doublet rate of 11.7%. In total, we generated 14,870 quality-filtered single-nucleus data sets having a median of 11,889 unique reads per cell that, from comparison with merged datasets, were confirmed to detect accessible regions (**Fig. S9-10**). To identify epigenetically-distinct subpopulations within this scTHS-seq dataset, we first used an unbiased approach. We modeled the probability of observing reads from a genomic site in a given cell as a censored Poisson process (see **Methods**), which accounts for the fact that the scTHS-seq signal from even the most accessible site will saturate after only a few reads. This approach revealed multiple distinct subpopulations in both the visual and frontal cortex (**Fig. 3B-D, Fig. S11**), with proportions that are more favorable to non-neuronal cell types than snDrop-seq **(Fig. 2E)**, suggesting that snDrop-Seq is less efficient in sampling non-neuronal cells, either due to data quality filtering that is biased against cells transcribing fewer transcripts or lower efficiency in packaging smaller nuclei into droplets. Characterizing the identity of epigenetically-defined subpopulations, however, is more challenging than in the case of transcriptionally-defined subsets, as functional roles of most regulatory sequences remains poorly annotated. Based on the functional annotation of the genes neighboring differentially-accessible sites, we could distinguish five glial cell populations and three neuronal cell populations having 97,672 differential accessibility peaks (**Table S8**) that when aggregated into accessibility profiles could be used to identify putative regulatory regions within a given locus (**Fig. 3D**).

**Fig. 3.**
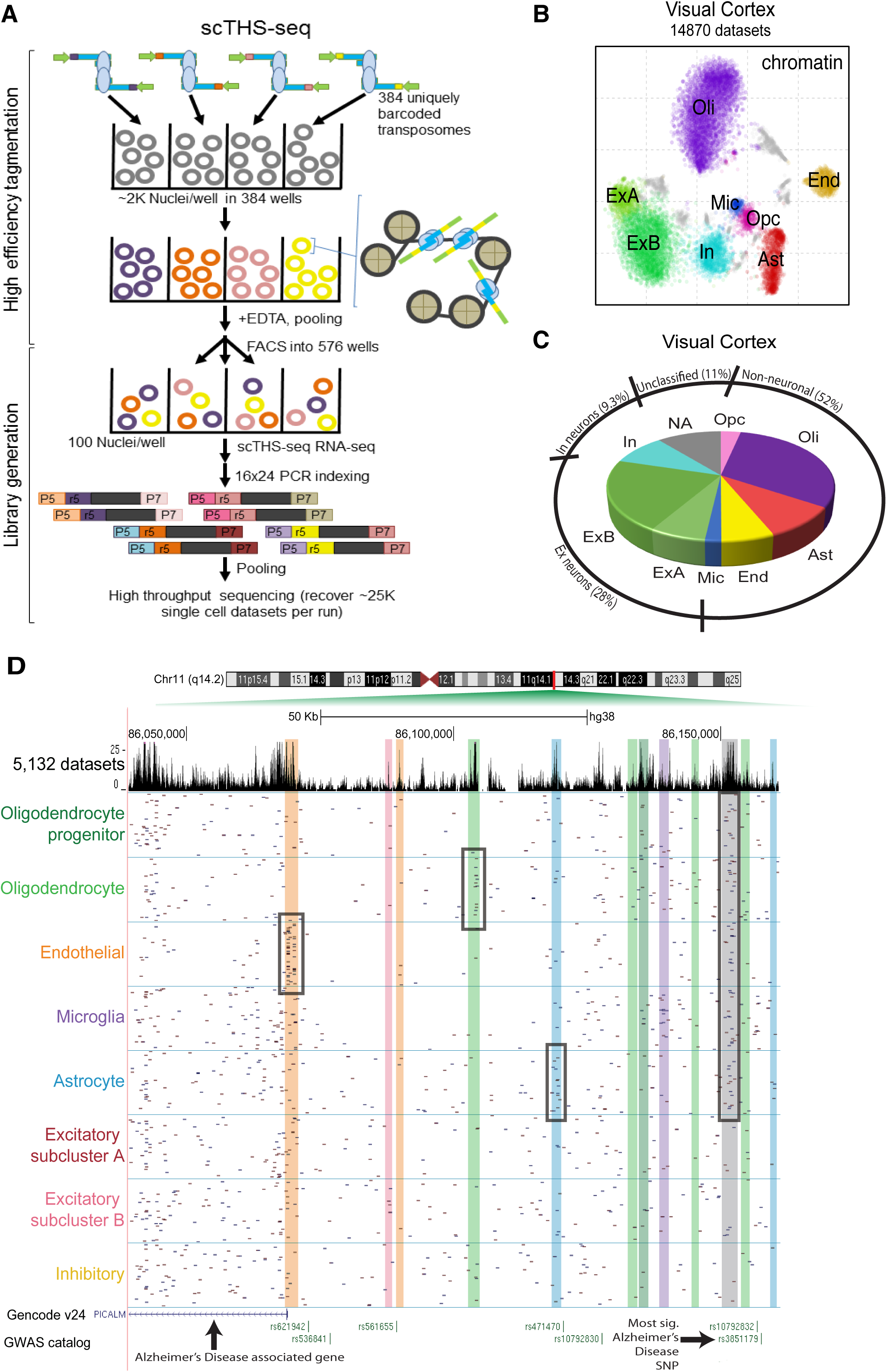
Chromatin accessibility profiling resolves major cell types of the visual cortex. **A.** Implementation of scTHS-seq with a Tn5 supermutant and cellular combinatorial indexing. First, 384 uniquely barcoded transposomes are added to 500,000 to 1 million mixed mouse and human nuclei that have been aliquoted into a 384 well plate, with one uniquely barcoded transposon added per well. Species nuclei are mixed to reduce and calculate collision rates, and for quality control. Next, high efficiency tagmentation is performed, followed by EDTA inactivation, pooling, and FACS redistribution of 100 nuclei/well into 96 well plates. scTHS-seq RNA-seq is performed with each well treated as one reaction, followed by PCR indexing, pooling and high throughput sequencing of libraries. Optimally ∼25,000 total single cell datasets are recovered per run, the sum of mouse and human nuclei recovered. **B.** tSNE plot of major subpopulations of visual cortex nuclei, clustered based on the overall similarity of the chromatin accessibility profiles. **C.** Pie chart depicting major visual cortex subpopulations identified in B. **D.** Identification of cell specific DNA accessibility peaks over the promoter and regulatory region of Alzheimer’s disease associated gene PICALM. Glial specific peaks are over-represented. The top tract is all cells merged to generate peaks. Each cell subpopulation tract is represented by 100 randomly selected single cells of cells that had reads in the depicted region, where each row represents a single cell and each dot is a read. The color of highlighted peak regions corresponds to the cell type specificity of the peak, with each subpopulation tract title a specific color. Boxed regions highlight specific cell type specific peaks. The gray box highlights a glial cell specific peak. Nonspecific peaks are not highlighted.

To establish more precise correspondence between transcriptional and epigenetic states of different subpopulations, we sought to identify cells corresponding to transcriptional subpopulations in the scTHS-seq data as well as cells corresponding to epigenetic subpopulations in the snDrop-seq data. We trained a gradient boosting model (GBM) to predict differentially accessible genomic sites based on the differential expression patterns (**Fig. 4A, Fig. S12**) and a separate GBM to predict differential expression based on differential accessibility (**Fig. S12-13**) using features such as the distance of a site to a gene, degree of differential expression or accessibility of the site or gene, among others (see **Methods**). While the ability to predict differential expression or differential accessibility of any individual gene or site is limited (**Fig. S12B,E**), joint consideration of multiple genes or sites allows for confident cell type classifications (**Fig. 4B,C, Fig. S12C,F, Fig. S13B,C**). Applying such models to classify scTHS-seq cells based on the differentially expressed genes between major cell types, we were able to resolve astrocytes, oligodendrocytes, inhibitory and excitatory neurons with high sensitivity and precision in both the visual cortex (**Fig. 4B**) and frontal cortex (**Fig. S13D**). Similarly, applying such models to classify snDrop-seq cells based on the differentially accessible sites between major cell types, we were likewise able to resolve these cell types with high sensitivity and precision in both the visual cortex (**Fig. S13B,C**) and frontal cortex (**Fig. S13E**). Although layer 4 excitatory neurons (L4 = **Ex2-4**) were not distinguishable from layer 5 and 6 excitatory neurons (L5/6 = **Ex5-8**) from an unbiased analysis of scTHS-seq data alone (**ExB in Fig 3B**), integrating differential expression information from the higher resolution snDrop-seq data allowed us to identify relevant differentially accessible sites to further partition scTHS-seq clusters (**Fig. 4D**). Similarly, such integrated analysis allowed us to identify epigenetic differences relevant to inhibitory neuron subtypes that appear distinguished by their developmental origin from subcortical regions of the medial or lateral/caudal ganglionic eminences^7, 26, 27^ (**Fig. 4E**). Thus, despite lower intrinsic cell type resolution of accessibility data compared to transcription, computational integration of both scTHS-seq and snDrop-seq allowed us to reconstruct detailed epigenetic profiles of fine-grained cell types within the brain, enabling investigations of the regulatory processes active within each cell type.

**Fig. 4.**
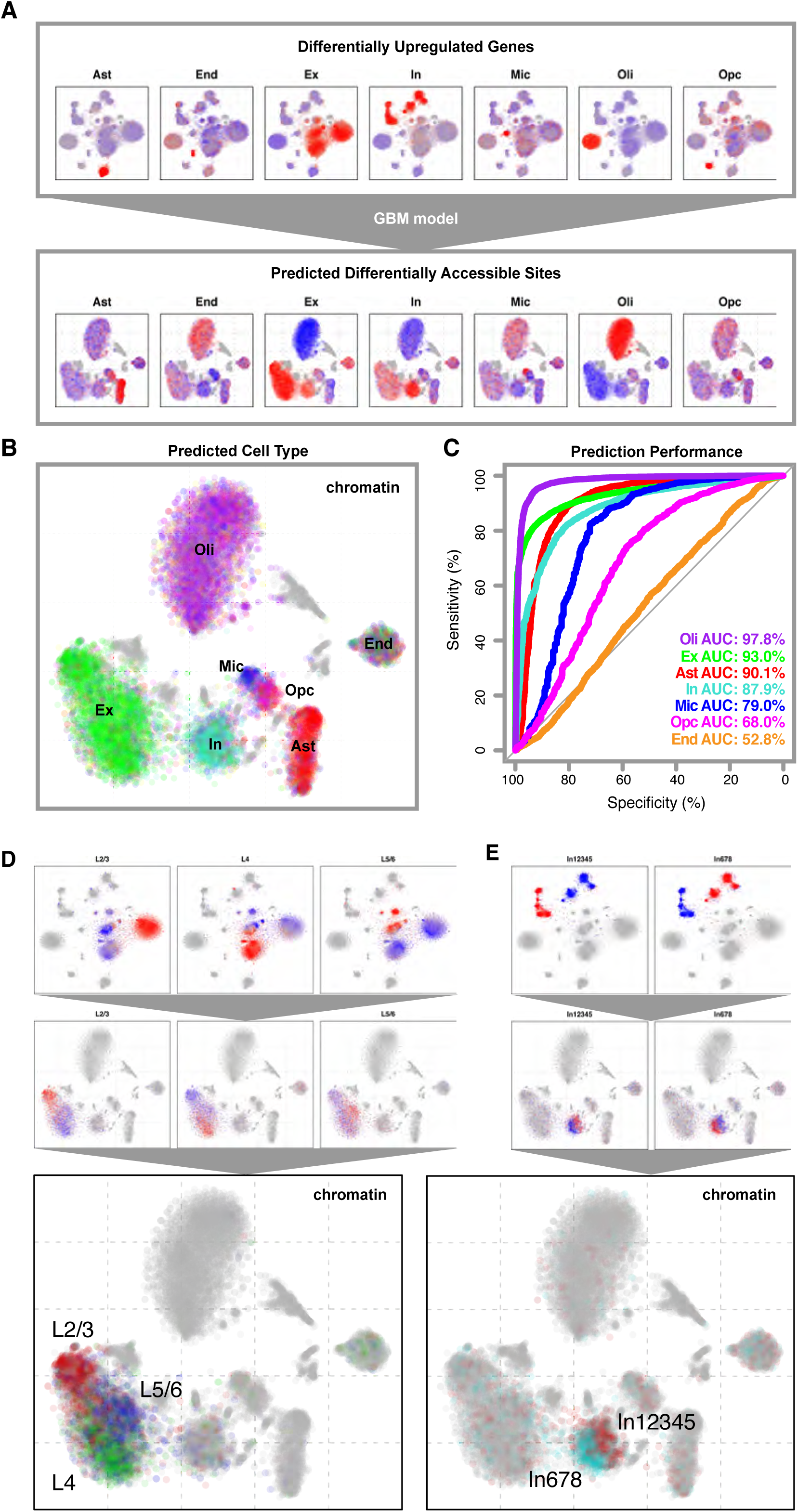
Differential gene expression predicts differential chromatin accessibility to map corresponding transcriptional and epigenetic subtypes. **A.** Average expression genes significantly upregulated (Z > 1.96) in each major cell type in the visual cortex are shown, with red corresponding to high expression of blue for low expression. A GBM model predicts differential accessibility of sites observed in scTHS-seq based on the predicted association of sites with the differentially expressed genes (see Methods). Predicted differentially accessible sites are visualized with red corresponding to high accessibility and blue for low accessibility. **B.** Predicted cell type identities are assigned based on the cell type with the maximal joint score on predicted differentially accessible sites. **C.** Prediction performance is measured by ROC curves and AUC comparing joint scores and previously inferred cell type identities identified from scTHS-seq analysis alone. **D.** Differential expression among excitatory neuronal subtypes predicts differentially accessible regions to further divide excitatory neuronal subtypes in the scTHS-seq data. **E.** Differential expression among inhibitory neuronal subtypes predicts differentially accessible regions to further divide inhibitory neuronal subtypes in the scTHS-seq data.

Having established the cell-type identity of each epigenetically-distinct subpopulation, we sought to identify the transcription factors (TFs) relevant to the regulatory state of each cell type. To do so, we looked for TFs whose predicted binding sites are over-represented within regions of differential chromatin accessibility distinguishing a given cell type. Screening a set of 379 TFs (**Table S9**) with position weight matrices from the JASPAR database^28^, we identified 195 TFs factors showing statistically significant (FDR <^10-5^) association with at least one of the cell types (**Fig. 5A-B, Table S10**). As expected, TFs associated with the neuronal subpopulations (**ExA**, **ExB**, and **In**) are distinct from TFs with binding relevant to non-neuronal subpopulations (**Oli**, **Ast, OPCs, Mic, and End**) (**Fig. 5B**). We also recover TFs with known relevance of cell subtypes. For instance *OLIG1*, an oligodendrocyte progenitor marker^29, 30^, showed enriched binding within the **Oli** cluster. As an independent validation, we integrated snDrop-seq data to confirm that the TFs showing significant association with a particular cell type also tend to show higher expression levels within that cell type (**Fig. S14**). Some of TFs achieve positive feedback by targeting their endogenous loci. For instance, *PKNOX2* has 8 of such binding sites within its own locus, 5 of which are differentially accessible, mostly in neuronal subpopulations compared to glia (**Fig. 5C**), highlighting the potential of this approach to identify positive self-regulation.

Cell-type specific epigenomics information has been highly valuable for identifying pathogenic cell types and specific regulatory mechanisms underlying many common genetic diseases^29-31^, yet brain diseases remained inadequately understood due to the lack of epigenomic maps with any cell-type resolution. To fill this gap, we obtained NIH GRASP database SNPs that were identified from genome-wide association studies (GWAS) as significant (p-values < 10^-6^) in ten brain related disorders, as well as seven additional diseases unrelated to the brain for controls. Given that causal variants are often located at different positions in linkage disequilibrium with the GWAS SNPs, we searched for enrichment of DNA accessibility regions in 100kb windows centered on all GWAS SNPs of a given disease, and assessed the significance by random permutations (**Fig 6A**, see **Methods**). This analysis identified strong enrichments in multiple cell types or sub-types, contrasting with an alternative possibility of uniformity (**Fig. 6B-C, Fig. S15, Table S13**), including: highly significant enrichment for common risk variants linked to Alzheimer’s Disease in **Mic** and **Oli** cells, Schizophrenia in **ExB** and **In** neurons, **Mic** and **Oli** cells; ALS in **In** and **Ast** cells; Parkinson’s Disease for the **ExB** and **In** neurons; Bipolar disorder for all **Ex** neurons and **Mic**; Autism for all **Ex** and **In** neurons; ADHD for al **Ex** and **In** neurons, **Oli** and **Opc**; ALS in **In** and **Ast** cells; with significant associations for Epilepsy for all **Ex** and **In** neurons and **Oli**, and for Depression for all **Ex** neurons and **Mic**. Significant enrichment for **Oli** was found in glaucoma, and similarly in the other non-brain diseases significant enrichments were not found in any neurons. However enrichments were found in the most closely related cell types to those implicated in the disease. For the autoimmune diseases Crohns, Celiac and Type I diabetes, **Mic** and **End** were enriched, while in lung disease **Mic** was enriched. No enrichment was found for the two non-brain related diseases (Chronic kidney disease and prostate cancer), demonstrating the specificity of these analyses. While further validation is required, our chromatin maps provide a new framework through which new aspects of brain diseases can be understood at the level of specific cell types or subtypes.

**Fig. 5.**
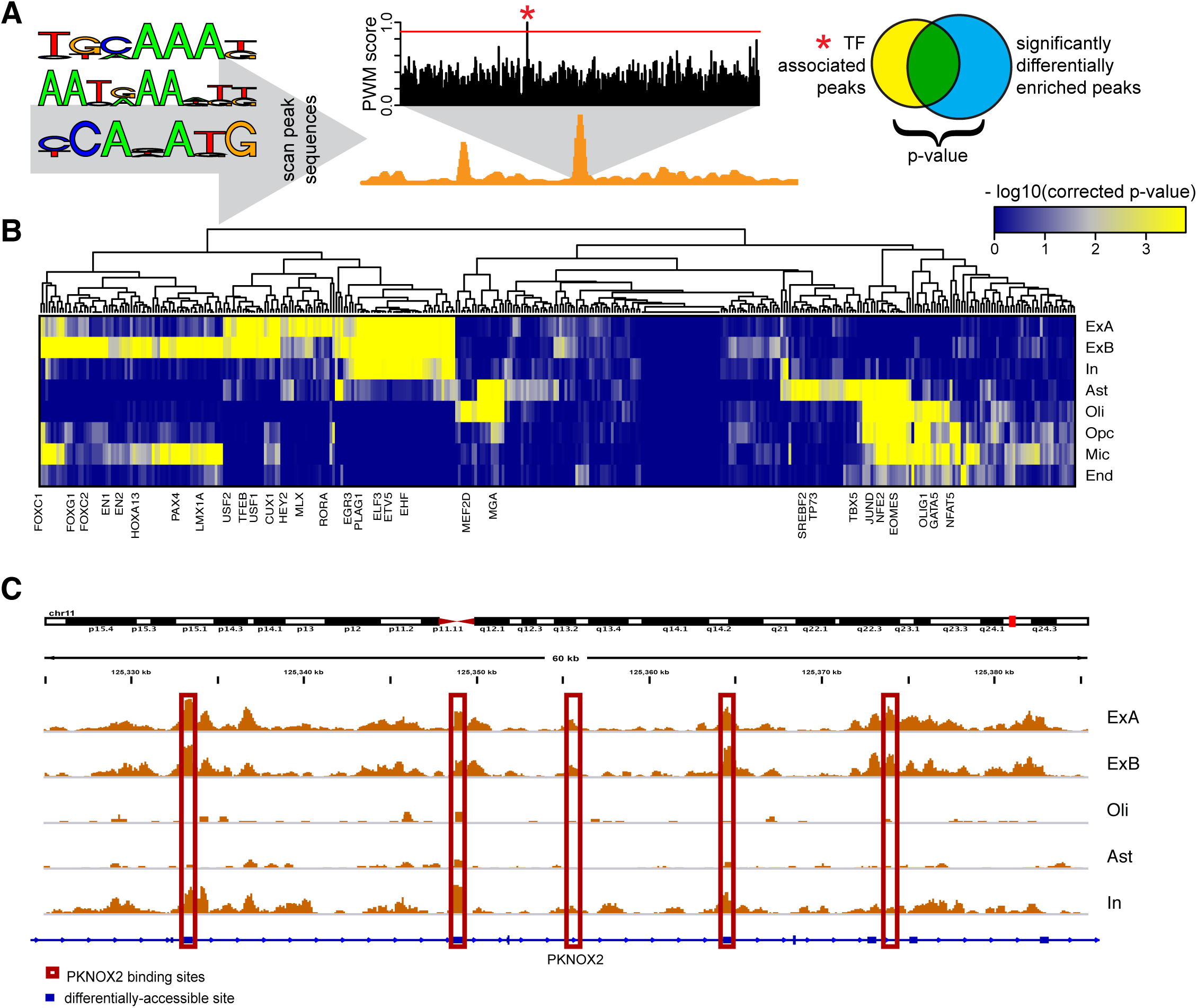
Mapping of transcription factor binding status and disease risk variants to specific brain cell types. **A.** Schematic of TF analysis. Briefly, putative TF binding sites (TFBS) were identified within all hypersensitive sites based on PWMs matching. To identify relevant factors for a given cell type, sites showing differential accessibility within that cell type were tested for statistical enrichment of different TFBS. **B.** Heatmap of TF association to epigenetic subpopulations. Each column is a TF. Each row is an epigenetic subpopulation. Select TFs are annotated. **C.** PKNOX2 transcription factor shows potential for self-regulation. The IGV view of a representative region for tracks corresponding to 5 subpopulations highlights patterns of differential accessibility in the PKNOX2 gene. Differentially-accessible sites within PKNOX2 are noted with blue boxes. Putative binding sites for PKNOX2 based on PWM sequence similarity within PKNOX2 are noted with red boxes.

**Fig. 6.**
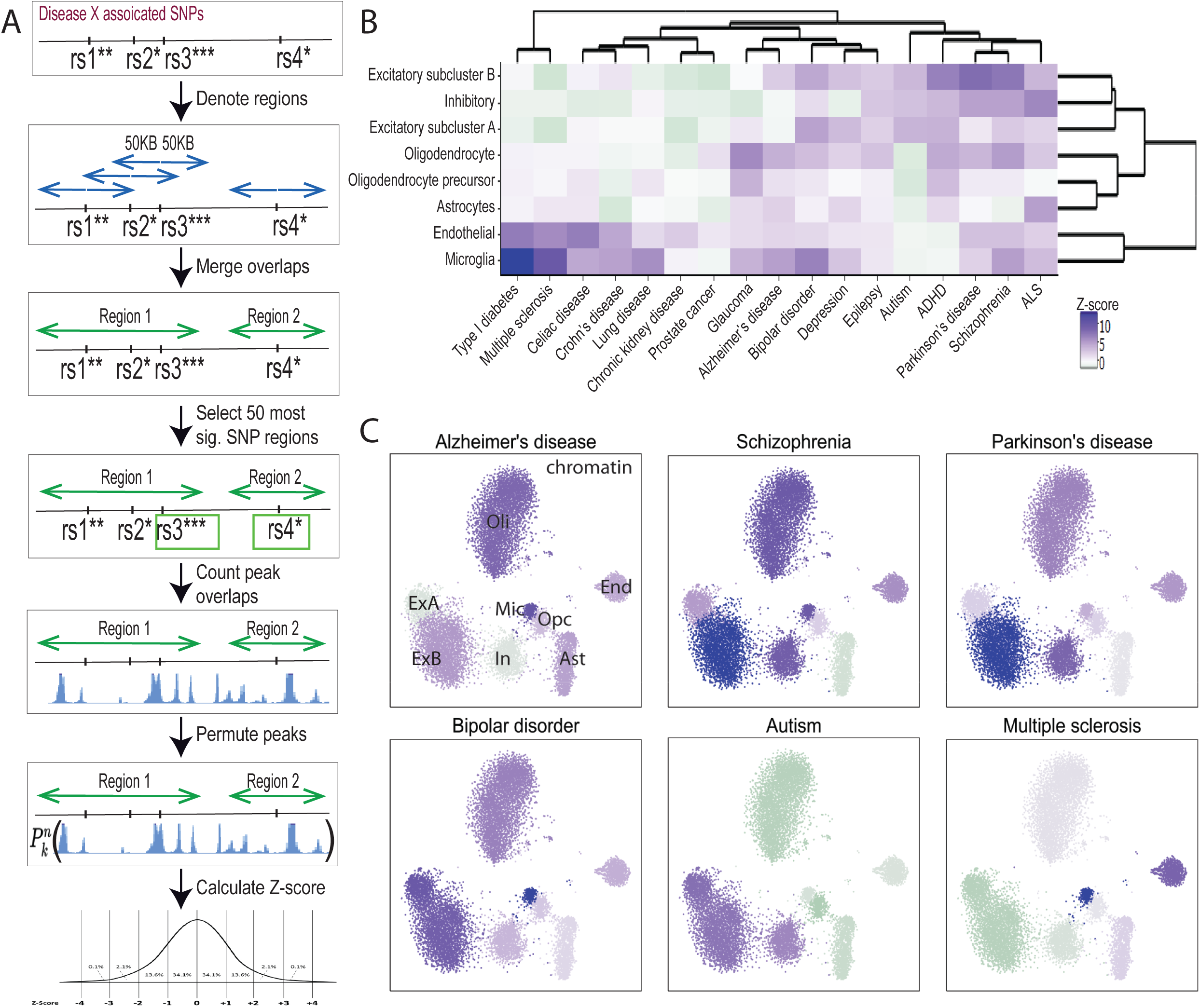
Mapping of common disease risk variants to specific brain cell types. **A.** Method used to map disease risk variants to specific cell types. Briefly, GWAS SNPs were obtained for each disease, extended to 100KB, merged, the top 50 most significant SNPs selected, number of peaks in overlaps counted, peaks permuted and the number of peaks counted in each region for each permutation, then lastly Z-scores were calculated. **B.** Heat map representing the enrichment Z-scores across 8 cell clusters (rows) for 10 brain diseases (columns) and 7 unrelated diseases. Dark purple and purple represent a significant Z-score over 1.96, where light purple, gray and light green represent an insignificant Z-score, and green represents a significant negative association with a Z-score less than -1.96. **C.** Z-scores for the enrichment of GWAS SNPs in the open chromatin of **ExB, In, ExA, Oli, Opc, Ast, End, Mic**, populations were overlaid onto the cell clusters. Six brain disorders are shown.

## Discussions

Reconstruction of cellular composition is an important aim towards understanding the human brain. Our study provides a first demonstration of an integrative single-cell analysis on the human adult brain, utilizing two highly scalable methods for acquiring transcriptional and epigenetic information from post-mortem tissues: snDrop-Seq and scTHS-seq. Using nuclei isolation to overcome challenges associated with processing of post-mortem tissues, we recovered all known non-neuronal and most neuronal subtypes in human adult cortex. These assays rely on a much shallower sampling of many more cells than our previous efforts, yet we were able to resolve these cell subpopulations and provide informative aggregate profiles for each subtype. Our results underscore the power of sparse sampling of single cells in complexed tissues at a massive scale: as long as the data from the single cells are informative enough for clustering and “virtual sorting” into different groups^32, 33^, they can be combined into aggregate profiles that are as rich as bulk sequencing of different cell populations.

We demonstrate a computational strategy for mapping between transcriptional and corresponding epigenetic states that can be used to reconstruct aggregate epigenetic profiles for fine-grained cell types. Such profiles provide valuable insights into the regulatory processes and elements shaping the identity of different cell types, as well as their relevance to human disease. While previous studies have identified pathogenic cell types for many human common diseases, our analysis enabled an assessment of the relative impacts by common genetic risk alleles to multiple cell types in an organ. It provides a coherent framework to consolidate previous findings, such as the relative contributions of glia, microglia and neurons to sporadic Alzheimer’s disease^34^. Such information is critical for identifying effective therapeutic targets. Generating multiple types of –omics maps from single cells *en mass* also enabled us to leverage the strength of each method to improve the confidence of cell type assignment, greatly enriching cell annotations. This combined approach thus represents a highly scalable strategy for systematic construction of atlases composed of single-cell data for human organs like the brain and eventually, for the full human body.

## Acknowledgments

Flow cytometry was performed both at the UCSD Human Embryonic Stem Cell Core and TSRI Flow Cytometry Core. We thank TF Osothprarop and MM He for providing Tn5059 transposase, N Salathia for assistance in sequencing, Y Wu for assistance in sequencing analysis. Funding support was from the NIH Common Fund Single Cell Analysis Program (1U01MH098977). TED was supported by a NIH K12 fellowship. GEK was additionally supported by Neuroplasticity of Aging Training Grant (5T32AG000216-24). PVK was supported by NIH Centers for Excellence in Genomic Science (P50MH106933) and NSF CAREER (NSF-14-532) awards. JF was supported by NIH grant F31 CA206236.

